# Coordinate based meta-analysis of whole-brain voxel-based morphometry studies does not show evidence of grey matter loss specific to PTSD

**DOI:** 10.1101/265496

**Authors:** Christopher R. Tench, Radu Tanasescu, Ketan D. Jethwa, Cris S. Constantinescu

## Abstract

Neuroimaging studies have detected structural alteration in post-traumatic stress disorder (PTSD), but findings are inconsistent. This might be explained by heterogeneity between subjects with PTSD in terms of common comorbidities such as depressive and anxiety disorders and also in traumatic experience. Despite this, coordinate based meta-analysis (CBMA) has been used to try and identify localised grey matter changes, and does suggest some PTSD specific pathology. However, there are multiple technical issues that make the meta-analytic evidence questionable, warranting a re-evaluation.

A literature search for voxel-based morphometry studies was performed. Only whole-brain studies using subjects with a current diagnosis of PTSD, and having a comparison group of either healthy or trauma exposed controls, were included. Twenty one voxel-based morphometry studies met the inclusion criteria. CBMA was performed to identify altered grey matter (GM) structures.

Using a novel coordinate based random effect size meta-analysis, no grey matter structure was identified as being consistently altered in PTSD compared to controls. This was also verified using the activation likelihood estimate algorithm.

There is no evidence, from CBMA, of consistent localised grey matter changes specific to PTSD. Inconsistency may reflect true heterogeneity in PTSD pathology or methodological issues with imaging and/or analysis, limiting the detection of PTSD specific pathology.

## Introduction

Post-traumatic stress disorder (PTSD) is a mental health condition characterised by symptoms of excessive fear response that can be reliably provoked. Regardless of trigger, PTSD causes clinically significant distress, and impairs social interaction and capacity to work. Neuroimaging has been used to explore structural brain differences between PTSD and control groups using voxel based morphometry (VBM), which compares either GM concentration or absolute volume [1]. These are essential to the understanding of the condition, and indeed neurocircuitry models of PTSD have been suggested [2] based, at least in part, on evidence gathered from neuroimaging [3]. The amygdala is involved with fear processing and hyperactivity of the amygdala is associated with PTSD. This is thought to be the result of impaired inhibitory activity from the ventral/medial prefrontal cortex (vmPFC) and hippocampus [2]. Evidence of reduced volume of these structures in PTSD patients has come from region of interest (ROI) based neuroimaging studies and has been summarised in meta-analyses [4, 5]. The functional relationship of the amygdala and the medial prefrontal cortex has also been found to be inversely correlated to symptom severity [6].

Neuroimaging experiments involving a-priori regions of interest have been performed in PTSD, but are inevitably prone to publication bias. VBM is less biased in that regions hypothesised to be affected can be analysed in context with whole brain effects; publication bias is still a concern, but not directed specifically at hypothesised brain structures. Reports of VBM experiments include the coordinates (in standardised stereotactic space) of abnormalities detected anywhere in the brain. To be of any significance beyond the study cohort the results must be at least partially PTSD specific. However, studies often involve small subject numbers, which can limit generalisability [2]. Furthermore, while within-study the subjects may have experienced the same traumatic event, between-studies there is heterogeneity. There is also heterogeneity between subjects in terms of common comorbidities such as depressive and anxiety disorders [7]. Consequently, assessment of generalizable PTSD specific pathology using neuroimaging is not straightforward.

Whole-brain (WB) VBM analyses lend themselves to a coordinate based meta-analysis (CBMA) approach. The aim of CBMA is to use the coordinates from experiments testing related hypotheses to find the anatomical regions where they cluster together representing agreement across studies; such agreement is an unlikely chance event, suggesting true pathology. Compared to VBM, where the resulting clusters represent voxel-wise significant grey matter differences, CBMA results generally relate only to the distribution of reported foci (coordinates) of the VBM clusters [8-10]. As with any meta-analysis (MA) the aim is to obtain better estimates of true effect, in this case localised changes in brain structure associated with PTSD, than provided by individual studies. Multiple analyses have been published and each indicates that, despite the inherent heterogeneity, there is observable PTSD specific pathology. Unfortunately, technical limitations of these studies render the results, and conclusions, questionable. Here a new coordinate based random effect size (CBRES) meta-analysis algorithm is used to identify which of the findings across VBM studies are consistently observed features of PTSD, while addressing the issues that limit the validity of previous analyses.

Four meta-analyses of VBM in PTSD have been published [11-14], each with technical limitations. The assumption of independence of the studies is important in MA [15] but is often violated by multiple experiments using the same subjects [16]. In CBMA correlated studies can bias the results by imposing apparent concordance; all four published meta-analyses included multiple studies as independent, but which shared subjects. One analysis [11] included a study of a-priori region of interest [17], rather than whole brain analysis, in violation of the CBMA methodological assumptions. Known software implementation issues [18] affected two of the analyses [11, 12]. Finally two studies [13, 14] employed the effect size signed differential mapping (ES-SDM) algorithm [19]. This method has a slightly different aim to others, attempting to estimate the voxel-wise grey matter difference from just the reported foci summaries. To achieve this, however, it employs no principled control of the type 1 error rate [20], making it difficult to interpret as a meta-analysis, which by definition requires robust statistics. It has been shown that if the included studies report false positive results, these will be echoed in the results [8]. Indeed the authors of the method recommend analysing only high quality studies [21], however multiple of the included VBM studies used liberal uncorrected p-value thresholding that likely resulted in false positive findings [20]. Consequently the results of these analyses are difficult to interpret.

A statistically robust re-evaluation of the evidence for generalisable structural alterations from WB studies of PTSD is warranted. The aim of this study is to use CBRES meta-analysis to explore consistent abnormal localised GM volumes or concentrations. Particular emphasis is placed on the analysis of independent WB data; where studies use common subjects, coordinates are pooled as suggested in [10, 22] to form independent sets. Differences due to experimental factors, such as using corrected/uncorrected statistical threshold or comparing PTSD subjects to healthy controls (HC) or trauma exposed controls (TEC), that may affect the MA are considered using a sensitive omnibus test of differences [23]. Furthermore, an algorithm (ClusterZ) employing a principled [20], and interpretable, threshold approach to control false positive results is used. Finally, CBMA studies rarely provide the data used for analysis making them difficult to verify or reproduce. Therefore, in line with recommendations for scientific data analysis reporting [24], the coordinates used in this study are provided to allow further evaluation/validation.

## Methods

### Ethical approval

No ethical approval was required for this study.

### Study search and inclusion criteria

A literature search was performed using the search terms: [(“post-traumatic stress disorder” OR “PTSD”) AND (“voxel based morphometry” OR “grey matter volume” OR “gray matter volume”)]. Studies included in previously published CBMAs of PTSD were also considered for inclusion.

Inclusion criteria were: studies involving adult subjects with a current PTSD diagnosis, the comparison of grey matter volume or concentration in PTSD to either HC or TEC, reported coordinates in either Talairach [25] or MNI [26] space. Studies that only performed ROI analyses violate the assumptions of the CBMA and were excluded; for studies performing both whole-brain and ROI analyses, only the whole-brain results were included.

### Effect of experimental factors

Meta-analysis combines results from experiments that, in principle, measure the same effect in order to improve estimates. In doing so it is necessary to check for heterogeneity among the included studies that might arise due to experimental factors that do not relate to the effect. In conventional MA this is possible by computation of an inconsistency measure [27]. Such methods are not available in CBMA, but a test of whether variance in the spatial distribution of reported coordinates is partly explained by experimental factors is possible (see for example [28]) using the omnibus test. The algorithm is detailed in [23], but briefly described here.

The algorithm tests the combined p-values (sum, over coordinates, of the log p-values) using a method analogous to Fisher's combined probability test; but without the assumption of independent, uniformly distributed, p-values. Dividing the experiments into two groups by an experimental factor allows a p-value to be computed for each coordinate using a permutation test of difference between groups [23]. The resulting test statistic is the combined p-values. Under the null hypothesis there is no difference in the spatial distribution of coordinates between the groups, so a similarly distributed combined p-value would be obtained for a random permutation of the grouping variable. To perform the omnibus test the null distribution of combined p-values is generated using many random permutations of the grouping variable. An extreme value of the test statistic, compared to this null distribution, provides evidence of a significant difference in coordinate pattern due to the experimental factor. Such differences indicate sources of heterogeneity mean the CBMA would need to be interpreted with caution.

### Coordinate based meta-analysis

The aim of CBMA is to identify statistically where the reported coordinates (foci of GM loss) from independent studies testing a common hypothesis are in spatial concordance. The results are clusters representing the spatial distribution of reported foci located where there is consensus across studies that there is a significant effect (for example grey matter loss or bold response). The anatomical locations of the clusters are then assumed to indicate pathology relevant to the common hypothesis. Statistical significance is determined using a null hypothesis that models lack of spatial concordance between studies by replacing the reported coordinates by coordinates drawn at random from grey matter image mask. It is then required that the observed coordinates agree spatially significantly better than random coordinates.

This study employs a new algorithm [8], ClusterZ, which also considers the Z scores or t-statistics commonly reported with each coordinate. Each reported Z score is first transformed into an effect size and variance that is independent of the number of subjects in the study. Then in each cluster formed by the spatial concordance of coordinates a conventional random effect size meta-analysis, or meta-regression, is performed. Interpretation of each cluster is as for conventional MA, and the use of standard visualisation and data-checking tools such as forest plots is possible. ClusterZ has multiple advantages over other algorithms. In particular by utilising the reported Z scores the p-values are based on an effect size rather than a density of reported coordinates, which might depend on the size of the structure affected. Furthermore, ClusterZ has an easy to interpret approach to principled [20] type 1 error rate control through the use of the false cluster discovery rate (FCDR). This is the proportion of the clusters declared significant that would be expected just by chance. Being based on the false discovery rate (FDR) method [29] this is less conservative than the cluster-wise family wise error (FWE) method recommended with the ALE algorithm, and avoids the issue of employing voxel-wise FDR that can lead to false positive results [30].

### Experimental procedure

#### Data extraction

Studies identified were scrutinised for: coordinate space used (MNI or Talairach), number of subjects, subject characteristics, and whether corrected statistical analysis was employed. Coordinates and Z scores from each study were extracted, and those in MNI space converted to Talairach [31]. Talairach labels were determined as detailed in [32]. Both GM volume increase and decrease were extracted; the convention used is that PTSD patients having increased GM volume relative to controls have Z>0, while relative GM loss is indicated by Z<0. Extracted data are checked by two authors (CRT and RT). In addition demographic and relevant clinical variables were also extracted and tabulated.

#### Analysis

Studies entered into the CBMA must be independent to avoid bias. Coordinates from multiple experiments in single studies using the same subjects were therefore pooled into single independent studies. To look for shared subjects across different studies the papers were scrutinised for similar cohort demographics. Furthermore, ClusterZ automatically indicates where different studies report occurrences of identical coordinates, which is unlikely for independent experiments, as a pointer to potentially correlated reports. Different studies sharing subjects were pooled to form a single independent study. Where there was doubt, the authors were contacted.

Difference in spatial distribution of coordinates due to experimental factors was tested for using the omnibus test. Factors considered were: use of corrected or uncorrected p-values, control group (TEC or HC), and treated or untreated PTSD subjects. A p-value of <0.05 was considered significant, and indicative of a source of heterogeneity.

Coordinate based random effect size meta-analysis was performed, using ClusterZ, on studies comparing PTSD subjects to trauma exposed controls or healthy controls both collectively and by control subject type. An FCDR of 0 05 was used, meaning only 5% of clusters declared significant were expected by chance. Post hoc analysis was performed to detect the next most significant clusters (FCDR>0.05) to make sure none were just missed; FCDR estimates are provided for these clusters.

## Results

### Search results

The preferred reporting items for systematic reviews and meta-analyses (PRISMA) [33] flow chart is given for both VBM studies in figure 1.

**Fig 1.**
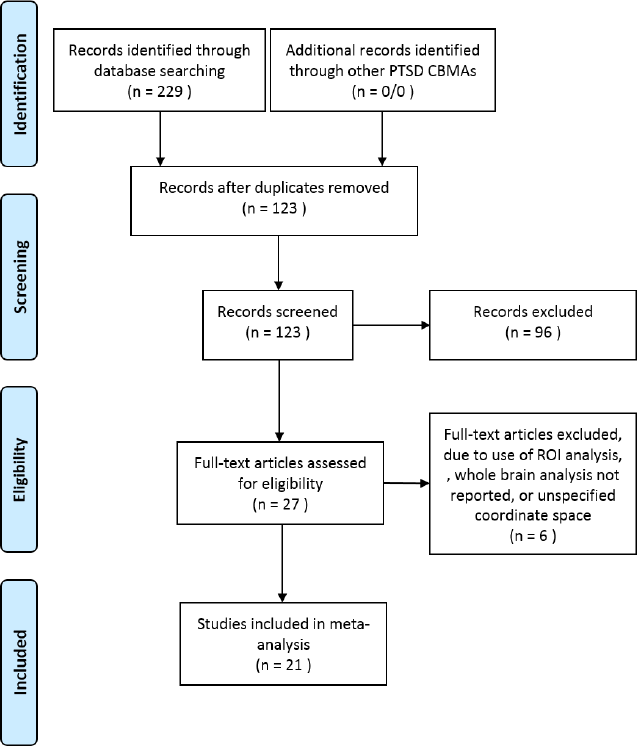
PRISMA flowchart depicting the study selection process for systematic reviews; numbers indicate VBM studies.

Fig 1. PRISMA flowchart depicting the study selection process for systematic reviews.

### VBM studies

Twenty one whole-brain VBM studies, reporting 24 relevant experiments involving a total of 168 coordinates, were suitable for inclusion; see table 1. These VBM studies overlapped substantially with those included in previous CBMAs, with some recent studies added [34, 35] and others omitted: Bryant et al [17], Eckhart et al [36], and Cortese et al [37] because they are ROI studies, Felmingham et al [38] because coordinates were only given for a-priori hypothesised regions, Hakamata et al [39] because the coordinates given fell outside of both Talairach and MNI space and no other coordinate space was specified, and Jatzko et al [40] because whole brain VBM results were not reported.

**Table 1.**
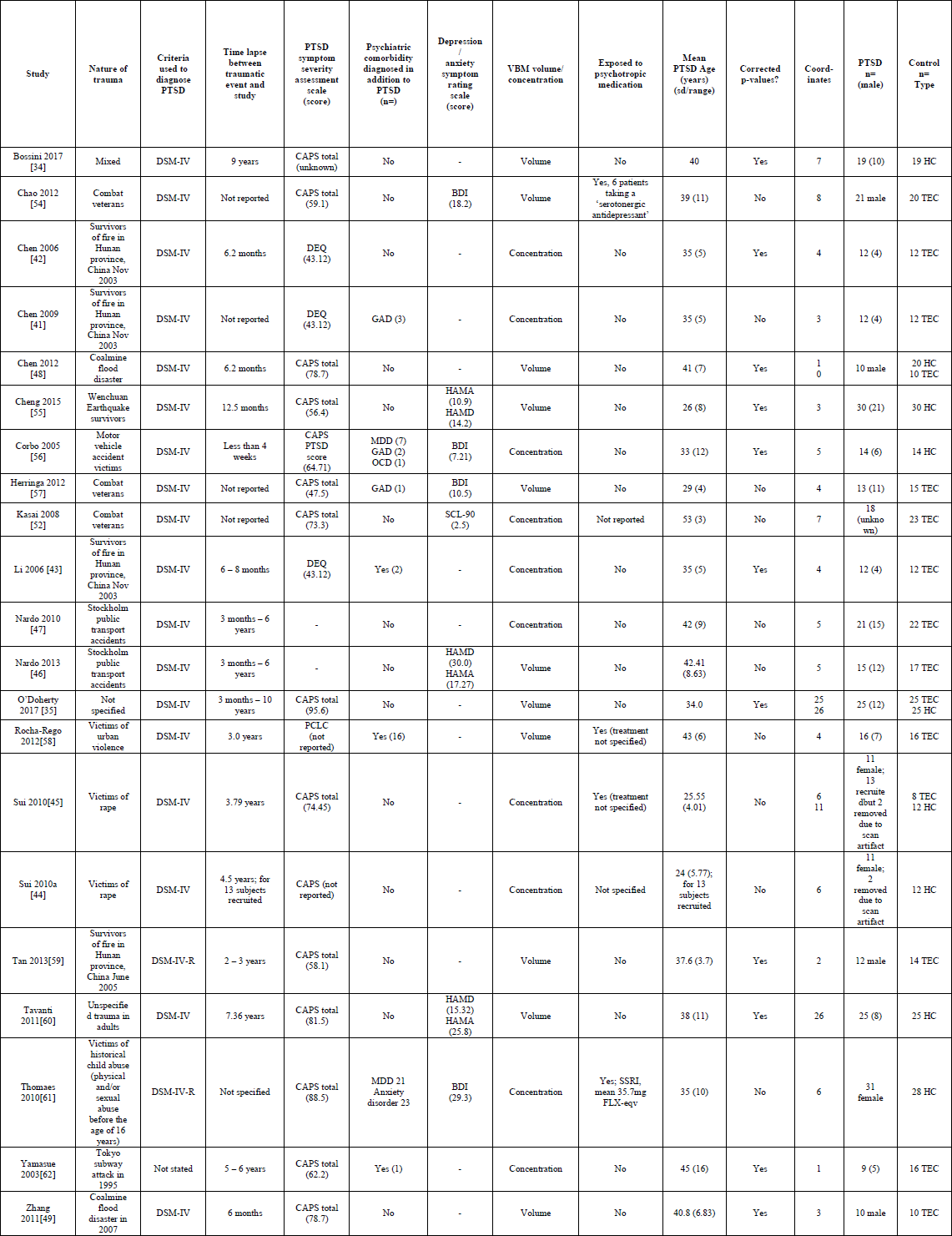
VBM studies included in analysis. Of these 21 studies, 24 experiments were extracted: 15 comparing PTSD to a TEC control group, and 9 comparing to a HC group. Principled methods of controlling the type 1 error rate were employed by 11 of the studies. Eleven of the studies used VBM to analyse GM volume, while ten analysed GM concentration. DSM: Diagnostic and Statistical Manual of Mental Disorders. CAPS: Clinician-Administered PTSD Scale. DEQ: Distressing Events Questionnaire. SCL-90: Symptom Check List-90. HAMA: Hamilton Depression Scale. HAMD: Hamilton Anxiety Scale. BDI: Beck Depression Inventory. GAD: Generalised anxiety disorder. OCD: obsessive compulsive disorder.

As noted by Li and colleagues [12], some of the studies used the same subjects. The survivors of a fire in the Hunan province of China were used in three of the studies [41-43], and two studies [44, 45] shared the same victim of rape subjects; the coordinates from these studies were merged such as to create two independent studies. The effect of pooling on significant clusters was also considered when subsets of subjects were potentially shared; the clusters should be robust against pooling of the data to ensure they are not driven by replicated results from the same subjects. The two studies by Nardo et al [46, 47] had some subjects in common (confirmed by author), and two studies potentially a subset of coal mine flood survivors [48, 49]. Taking into account these duplications: a total of (approximately, without complete knowledge of duplicated subject use) 316 PTSD subjects and 382 matched controls were studied.

### Effects of experimental factors

A comparison between VBM studies employing corrected and uncorrected p-values using the omnibus test detected no difference in the spatial distribution of coordinates (p=0⋅7). Similarly comparing VBM studies using healthy controls to VBM studies using trauma exposed controls also detected no difference in the spatial distribution of coordinates (p=0⋅6). Of those VBM studies that reported treatment status of the patients only four reported using a treated cohort, therefore no comparison was performed using the omnibus test.

From these comparisons there was no detectable evidence that the spatial distribution of coordinates was modified by the main experimental factors. The experiments were therefore considered homogeneous for the purpose of analysis.

### Meta-Analysis of VBM studies

On performing CBRES meta-analysis of all studies listed in table 1, no significant clusters were detected. The most significant cluster was not detectable below an FCDR of 0⋅34.

Results from the present analysis are contrary to previous CBMAs, which all reported multiple significant clusters. Therefore, independent validation was sought by performing the CBMA with GingerALE (version 2.3.6) using the recommended cluster based method method [50]. No significant clusters were detected in this analysis.

## Discussion

PTSD is a condition that is inhomogeneous in terms of comorbidities, and traumatic experience is certainly individual. It is perhaps of no surprise that the results of VBM studies are inconsistent. Despite this multiple coordinate based meta-analyses have suggested generalizable pathology specific to PTSD. Such findings are important as they can inform models of PTSD. Given the technical limitations of previously published CBMAs of PTSD, and that none provided data for validation, a re-evaluation of the neuroimaging evidence for consistently detectable neuropathology was warranted. Meta-analysis of the whole-brain VBM studies revealed no significant clusters, indicating a lack of detectable agreement across VBM studies of PTSD.

The results presented here contradict those from previously published CBMAs, despite the similarity of included studies. Explanations include software implementation issues, and methods employed to correct for multiple comparison. Other issues with previous CBMAs include: use of ROI studies that can bias the results as they are generally designed around neurocircuitry models of PTSD, and treating studies as independent when in fact they use the same subjects. However, it is not possible to know exactly what has caused this apparent contradiction without access to the coordinate database used; the coordinate databases used here are provided as supplementary material. To independently check that the results were reproducible, the same coordinates were analysed using the GingerALE software (version 2.3.6) and no clusters were detected.

Ideally the results of neuroimaging studies on PTSD would be generalizable. However, it is the nature of traumatic events that they often involve few people, so many studies use small (same traumatic event) sets of subjects. Lack of statistical power might then mean that real effects are simply missed; false negatives. To counter this liberal, uncorrected, statistical thresholds are used by many (see table 1), but this risks false positive results; indeed, one author noted that application of type 1 error control by correction for multiple comparisons completely eliminated all significant results [52]. The heterogeneity (different traumatic experience) of subjects between studies is another candidate explanation, since the specifics of the trauma are relevant to PTSD [53]. Heterogeneity in terms of experimental factors, such as comparison to a healthy or trauma exposed control group, might contribute to the overall lack of detected consistency, but this was not detectable using the omnibus test. Elapsed time since trauma (duration of disease) is another factor that is heterogeneous between studies. Administration of rating scales to assess affective symptoms in the included studies revealed that anxiety and depressive symptoms are common, and some studies included patients fulfilling the diagnostic criteria for a co-morbid depressive disorder. This reduces the diagnostic purity of the cohort and introduces a confounding factor. Past or present pharmacological treatment may also have an effect on imaging findings. Wherever possible, a sensitive omnibus test was employed to explore whether differences in experimental design had resulted in heterogeneity, but none were found to explain a significant amount of the variance in reported coordinates.

## Summary and Conclusions

The present coordinate based random effect size meta-analysis of voxel-based morphometry studies produced no evidence of consistent neuroimaging detectable structural abnormality in PTSD. The lack of results compared to previous coordinate based meta-analyses of PTSD is attributed to: software issues in previous studies, careful consideration for the independence of the studies included in the analysis, and explicit exclusion of results from a-priori region of interest analyses. These results suggest that either neuroimaging experiments of PTSD have a level of reproducibility that is not detectable by CBMA, or that pathology in PTSD is heterogeneous.

